# A generalizable computational mechanism underlying the interaction between momentary craving and decision-making

**DOI:** 10.1101/2023.04.24.538109

**Authors:** Kaustubh R. Kulkarni, Laura A. Berner, Shawn A. Rhoads, Vincenzo G. Fiore, Daniela Schiller, Xiaosi Gu

## Abstract

Substance craving and maladaptive choices are intertwined across addictive disorders. However, the computational mechanisms connecting craving and decision-making remain elusive. Here, we tested a hypothesis that momentary craving and value-based decision-making influence each other during substance-related reinforcement learning. We measured momentary craving as two groups of human participants (alcohol drinkers and cannabis users; total *n*=132) performed a reinforcement learning task in which they received group-specific addictive cue or monetary rewards. Using computational modeling, we found that, across both groups, momentary craving biased learning rate related to substance-associated prediction errors (RPEs), but not monetary RPEs. Additionally, expected values and RPEs jointly influenced elicited craving across reward types and participant groups. Alcohol and cannabis users also differed in the extent to which their craving and decision-making influenced each other, suggesting important computational divergence between the two groups. Finally, regressions incorporating model-derived parameters best predicted substance use severity in the alcohol, but not cannabis group, supporting the utility of using these model-based parameters in making clinical predictions for selective substance groups. Together, these findings provide a computational mechanism for the interaction between substance craving and maladaptive choices that is generalizable across addictive domains.

## INTRODUCTION

Humans can become addicted to a variety of substances, including a wide range of drugs and alcohol. Two elements are essential across all types of addictions: *craving*, the strong subjective desire for a substance; and *decision-making*, the objective choices made by affected individuals. While it is universally acknowledged that craving and decision-making are tightly intertwined in addiction, the computational mechanism underlying this interaction is not clear. Historically, the cue-reactivity literature – one of the most influential characterizations of craving – emphasizes the elicitation of craving in response to learned addictive cues that serve as secondary drug rewards^1–7^. Cue reactivity paradigms have been widely used to identify the neural correlates of craving (e.g., midbrain, insula, and cingulate) across a number of addictive disorders and sensory modalities^8–11^. Yet, they do not provide a mechanistic explaination for how craving arises in or interacts with drug-related choice behaviors. For example, forced abstinence and associated removal of drug cues paradoxically leads to increased craving and drug-seeking behaviors in substance dependent rodents^12,13^ and humans^14,15^, a phenomenon termed incubation of craving. Furthermore, it has been shown that drug-related beliefs and expectations also affect craving, an effect independent from the availability of drug rewards and cues^16–18^. As such, despite a rich empirical literature on craving, the computational mechanisms of craving remain elusive.

Computationally, reinforcement learning (RL) has been a primary framework used to account for maladaptive choices in addiction, with a central tenet that choices are reinforced by reward prediction errors (RPEs). Preliminary computational models hypothesized that addictive stimuli produce an irreducible RPE signal, subserved by excessive dopamine, that continuously reinforces substance-related choices^19^, which then subsequently shift one’s homeostatic setpoints^20^. While views on heterogeneity of dopaminergic encoding of information have become more nuanced^21–23^, and modern accounts have provided compelling evidence for RL-based behavior in animal models of addiction, these theories have yet to be tested empirically in humans with substance dependence. Critically, they also still do not account for how drug-related choices and craving may mutually influence each other. Recent efforts in computational psychiatry have started to shed light into the interaction between value-based decision-making and subjective states such as mood^24–29^. For example, monetary RPEs were found to predict mood ratings, providing initial evidence that internal subjective states could be influenced by RL signals in a systematic fashion^28^. Conversely, mood may increase the ‘momentum’ of value updating, providing a plausible mechanism for how mood drives dynamic changes in learning^24^. Momentary craving is likely entangled with addictive decision-making in similar ways, yet only a handful of empirical studies have examined this important relationship^16,30^, and a computational mechanism linking these two constructs remains noticeably missing despite well-established theoretical and empirical accounts of addiction that connect the two^19,31–33^.

In this study, we tested the hypothesis that momentary substance craving and value-based decision-making shape each other in a bidirectional fashion in humans. To test this hypothesis, we developed a paradigm in which substance-using individuals made choices to obtain either monetary or addictive cue (i.e., alcohol or cannabis) outcomes, and intermittently self-reported their craving (i.e., for alcohol or cannabis) during both blocks. To test the generalizability of our hypothesis, we examined two groups of participants (total n=132; see **Table 1** for participant characteristics): alcohol drinkers (n=68), and cannabis users (n=65). The task consisted of a modified two-armed bandit (**Fig. 1a**), where participants selected one of two machines (80% reward rate), and saw the outcome of either a coin (in the money condition) or their pre-selected addictive cue of either alcohol or cannabis (in the addictive condition). Momentary craving and mood were both sampled during the task (33% and 20% of the trials, respectively), and a novel computational modeling approach was used to fit both choice and craving data per session. We found that, across both groups, momentary craving biased *learning rate* in the addictive context, but biased *reward perception* in the monetary context. Conversely, in both substance and monetary contexts, elicited craving was driven by prediction errors (RPEs) *and* expected values (EVs), as opposed to either alone. Finally, we found that computational parameters derived from our models provided greater power than model-agnostic metrics for predicting clinical severity scores for the alcohol group, whereas demographics better predicted cannabis use severity. Together, these results validate a generalizable computational mechanism linking momentary craving with value-based decision-making in addictive disorders.

**Fig. 1.**
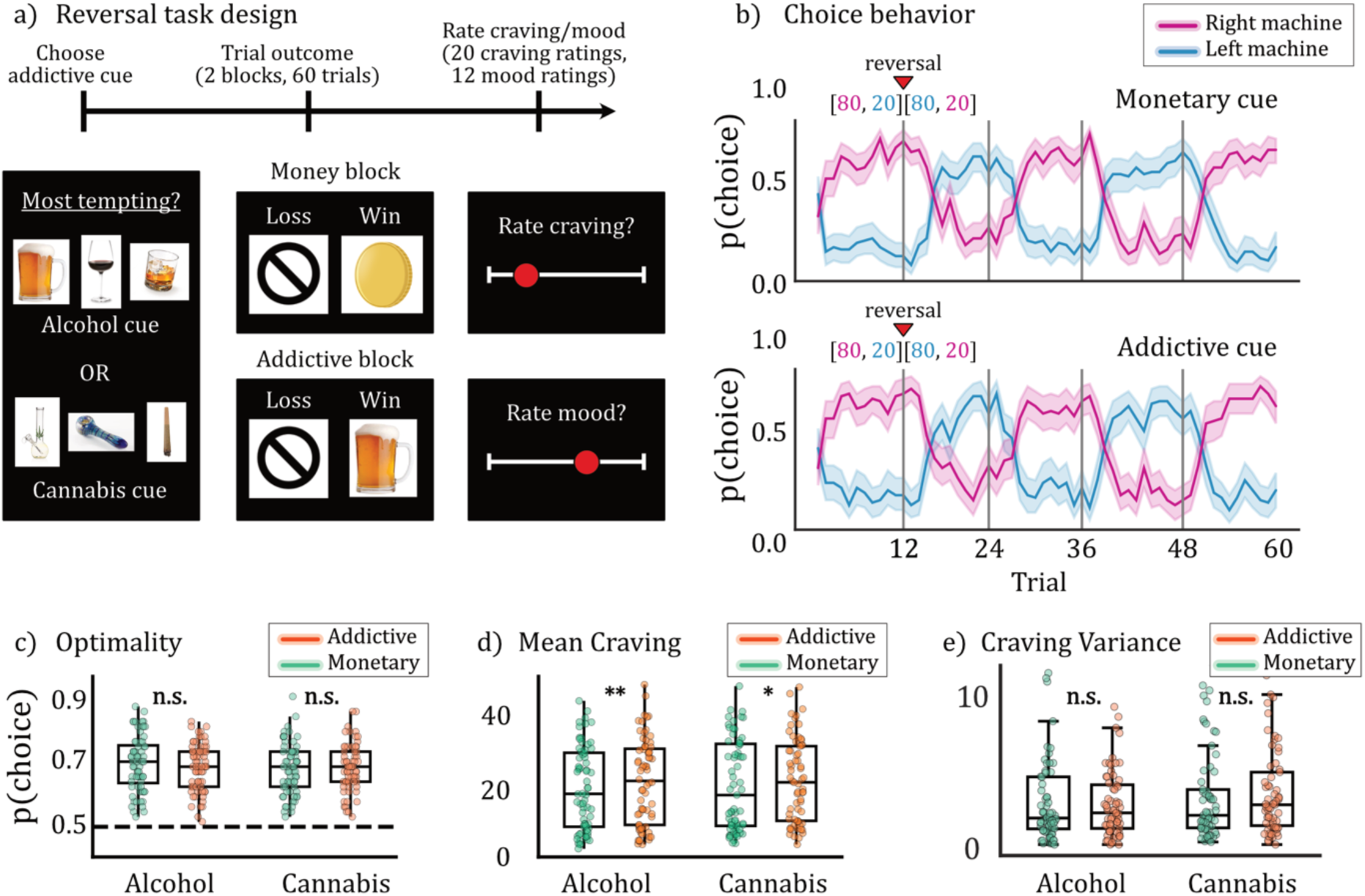
Experimental paradigm and model-agnostic task behavior. **(a)** Participants played a modified two-armed bandit task. Each session started with instructions and a choice of three reward options, of which the participant selected the one most tempting to them. Following five practice trials, two blocks of sixty trials were presented, in which the participant selected one of two machines. In the money block, the reward presented was an image of a coin. In the addictive cue block, the reward presented was the option selected by the participant at the start. Craving and mood were intermittently assessed throughout the block (20 craving ratings, 12 mood ratings), scored from 1 to 50. The more rewarding machine in each trial rewarded with 80% probability while the less rewarding machine rewarded at 20% probability. The optimal machine switched a total of 4 times (4 reversals) over the course of each block. The order and timings of the reversals was pseudorandomized over the cohort. Choice optimality was measured by calculating the proportion of participant choices that matched the optimal choice at each trial (i.e., the machine that rewarded 80% of the time at that trial). **(b)** Choice optimality was assessed visually by averaging participant choices across the experiment. Blue lines represent the left machine, purples lines represent the right machine, and vertical gray lines represent the timings of reversals in reward structure. In both groups, participants were able to successfully learn the reversal structure, reflected in the periodic switches of left/right machine choices over the experiment. **(c)** Choice optimality was calculated for each participant and stratified by group and condition. In both groups, choice optimality was significantly higher than chance (50%) (alcohol: *t*=18.9, *P*<0.001; cannabis: *t*=18.2, *P*<0.001). **(d)** Average craving ratings were significantly higher in the addictive cue condition compared to money condition in both groups (alcohol: *t*=3.14, *P*=0.002; cannabis: *t*=2.47, *P*=0.016). (e) Variances in craving ratings were not significantly different between conditions for either group (alcohol: *t*=0.793, *P*=0.431; cannabis: *t*=0.841, *P*=0.403).

**Table 1.** Participant characteristics.

| **Variable** | **Alcohol group**<br>**(n=68)** | **Cannabis group**<br>**(n=64)** |
| --- | --- | --- |
| Age (s.d.) | 40.9 (13.4) | 38.7 (14.1) |
| Sex (%) |  |  |
| Male | 45 (66.1%) | 37 (57.8%) |
| Female | 22 (32.4%) | 27 (42.2%) |
| Other | 1 (1.5%) | 0 (0%) |
| Education level (%) |  |  |
| ≤ High school | 10 (14.7%) | 7 (10.9%) |
| College | 42 (61.8%) | 44 (68.8%) |
| ≥ Graduate | 16 (23.5%) | 13 (20.3%) |
| Race (%) |  |  |
| American Indian or Alaska Native | 2 (2.9%) | 2 (3.1%) |
| Asian | 7 (10.3%) | 7 (10.9%) |
| Black/African American | 5 (7.4%) | 2 (3.1%) |
| Hispanic or Latino | 1 (1.5%) | 4 (6.3%) |
| White | 53 (77.9%) | 49 (76.6%) |
| ASSIST (s.d.) | 17.1 (8.4) | 14.0 (7.9) |
| AUDIT (s.d.) | 6.8 (6.6) | - |
| CAST (s.d.) | - | 7.7 (3.6) |

ASSIST – Alcohol, Smoking and Substance Involvement Screening Test<sup>87</sup>
AUDIT – Alcohol Use Disorders Identification Test<sup>90</sup>
CAST – Cannabis Abuse Screening Test<sup>92</sup>

## RESULTS

### Participants learned to maximize outcomes and reported fluctuating levels of craving

First, we examined model-agnostic behaviors that reflected how participants performed the task. In typical monetary bandit tasks, it is well established that humans learn to choose the option that maximizes monetary outcomes^34–36^; yet it remains unclear if individuals with moderate to heavy substance use behave similarly in the addictive stimulus condition. As in standard monetary tasks, here we defined choice optimality as the percentage of choosing the correct option (i.e. choosing the machine with the higher reward rate of 80%). Overall, participant choices were highly similar to the true reversal structure of the task (**Fig. 1b**), and choice optimality was significantly higher than chance (50%) regardless of condition (**Fig. 1c**; alcohol/money: 71±9%, alcohol/addictive: 70±8%, cannabis/money: 69±8%, cannabis/addictive: 70±8%, all *P*<0.001), confirming that participants successfully learned to exploit the machines for both addictive and monetary rewards. Choice optimality did not differ across participant groups (*t*=1.01, *P*=0.31), task conditions (*t*=0.38, *P*=0.71), or show an interaction effect (*F*=0.44, *P*=0.51), ensuring that findings related to craving-choice computation would not be attributable to distinct task performance alone.

Next, we verified that that participants experienced dynamic changes in their substance craving during the task, by calculating the mean and variability in self-reported cravings during the task across groups and conditions. As predicted, substance craving was greater in the addictive than the monetary condition in both groups (**Fig. 1d**; alcohol: *t*=3.141, *P*=0.002; cannabis: *t*=2.465, *P*=0.016), suggesting that addictive cues increased craving in this task, replicating cue-induced effects on craving. We also observed substantial variability in craving ratings within-subjects (**Fig. 1e**) such that craving variances were greater than zero for across groups and conditions (all groups and conditions; *t*>7, *P*<0.001), validating engagement during self-report of craving and dynamic changes in perceived craving in response to outcomes. Variances did not differ by group (*F*=0.26, *P*=0.61), condition (*F*=1.07, *P*=0.30), or their interaction (*F*=0.17, *P*=0.68). Finally, we also examined whether participants’ craving ratings might be inherantly correlated with mood, as negative affect has been found to be associated with increased drug craving^37,38^. To assess this, participant mood ratings were measured intermitently (“what is your mood right now?”) for 20% of the trials. Within-participant craving and mood ratings were not significantly correlated in the money condition (**Supp. Fig. 1**; mean correlations; alcohol: *r*=-0.06, *P*=0.31; cannabis:*r*=-0.002, *P*=0.96), or in the addictive cue condition (alcohol: *r*=-0.02, *P*=0.70; cannabis:*r*=0.06, *P*=0.31), indicating that craving ratings contained distinct information from mood ratings.

### A generalizable computational mechanism linking momentary craving and decision-making

Next, we constructed computational models that represent the bidirectional relationship between momentary craving and choice behavior (see **Methods; Table 2** for details). First, we composed five candidate model classes to account for choice behavior, with a modulation parameter (φ) defining the degree to which momentary craving modulated different components of the decision process (TDRL (temporal difference reinforcement learning, no bias)), Reward bias (r-bias), Learning rate bias (*α*-bias), Temperature bias (*β*-bias), and Momentum-based TDLR (m-TDRL)). The first four models were derived from classic TDRL models^39,40^ from the RL literature. In the r-bias model, momentary craving modulated the perceived magnitude of the reward signal^41,42^, while it instead modulated the learning rate and softmax temperature parameters in the *α*-bias and *β*-bias respectively. Finally, the m-TDRL model conceptualized craving as “momentum”, similar to recent efforts in modeling mood dynamics^24,43^.

**Table 2.**
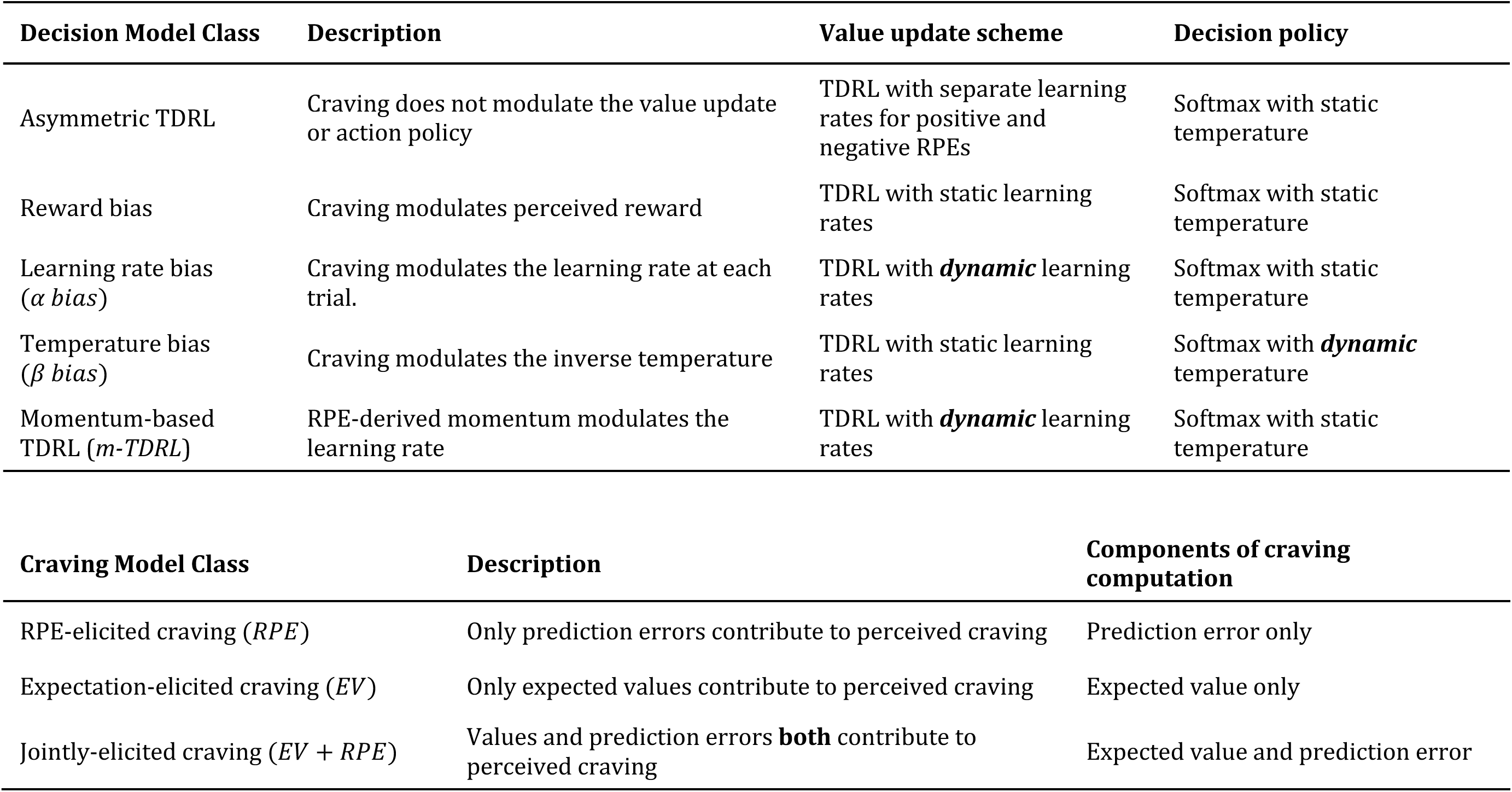
Decision and craving models.

| Decision Model Class | Description | Value update scheme | Decision policy |
| --- | --- | --- | --- |
| Asymmetric TDRL | Craving does not modulate the value update or action policy | TDRL with separate learning rates for positive and negative RPEs | Softmax with static temperature |
| Reward bias | Craving modulates perceived reward | TDRL with static learning rates | Softmax with static temperature |
| Learning rate bias ($\alpha$ bias) | Craving modulates the learning rate at each trial. | TDRL with *dynamic* learning rates | Softmax with static temperature |
| Temperature bias ($\beta$ bias) | Craving modulates the inverse temperature | TDRL with static learning rates | Softmax with *dynamic* temperature |
| Momentum-based TDRL ($m$-TDRL) | RPE-derived momentum modulates the learning rate | TDRL with *dynamic* learning rates | Softmax with static temperature |

| Craving Model Class | Description | Components of craving computation |
| --- | --- | --- |
| RPE-elicited craving ($RPE$) | Only prediction errors contribute to perceived craving | Prediction error only |
| Expectation-elicited craving ($EV$) | Only expected values contribute to perceived craving | Expected value only |
| Jointly-elicited craving ($EV + RPE$) | Values and prediction errors **both** contribute to perceived craving | Expected value and prediction error |

Next, we constructed models where different components of decision variables and their combinations contributed to future craving ratings. These models were inspired by 1) a recently proposed theoretical framework that craving arises as a posterior inference stemming from both prior expectations and prediction errors generated by outcomes^44^; and 2) computational models of other types of subjective states such as mood^28^. Three model classes were constructed and compared: RPE-elicited models (where only prediction errors influced craving), EV-elicited models (where only expected values influenced craving) and a full model with both RPE- and expectation-elicited craving (jointly-elicited craving). Note that in our specification, RPE-elicited craving essentially represents classic cue-induced craving because outcomes and RPEs are highly correlated (**Supp. Fig. 2**), while EV-elicted craving represents a testable alternative that prior beliefs are important in eliciting craving, a hypothesis that has gained increasing empirical validation in recent years^16,30,45^.

### Momentary craving biases drug-related learning across cannabis and alcohol groups

Of our candidate models linking craving and valuation, which best explains the choices made by participants? Following model comparison (**Fig. 2a, b**), we found that the *α*-bias model performed best in the addictive-cue condition across both alcohol- and cannabis-using groups. To assess the fidelity of fit by this model, we generated 2,000 simulations of choice behavior and calculated the degree of alignment between simulations and true behavior. We found that simulated behavior matched true behavior significantly better than chance (**Supp. Fig. 3**, both conditions: t>10, *P*<0.001) and close to optimal (**Supp. Fig. 4**), and parameter recovery was excellent (**Fig. 2c, d**). Examination of the parameter values for this model (**Fig. 2e, f**) revealed that φ was positive in alcohol users (M=0.209, SD=0.798, *P*=0.034) and negative in cannabis users (M=-0.995, SD=1.435, *P*<0.001), suggesting that higher craving accelerated alcohol-related learning for alcohol drinkers but slowed down cannabis-related learning for cannabis users. In other words, alcohol craving increases one’s sensitivity towards alcohol-related prediction errors, whereas cannabis craving shows the opposite effect.

**Fig. 2.**
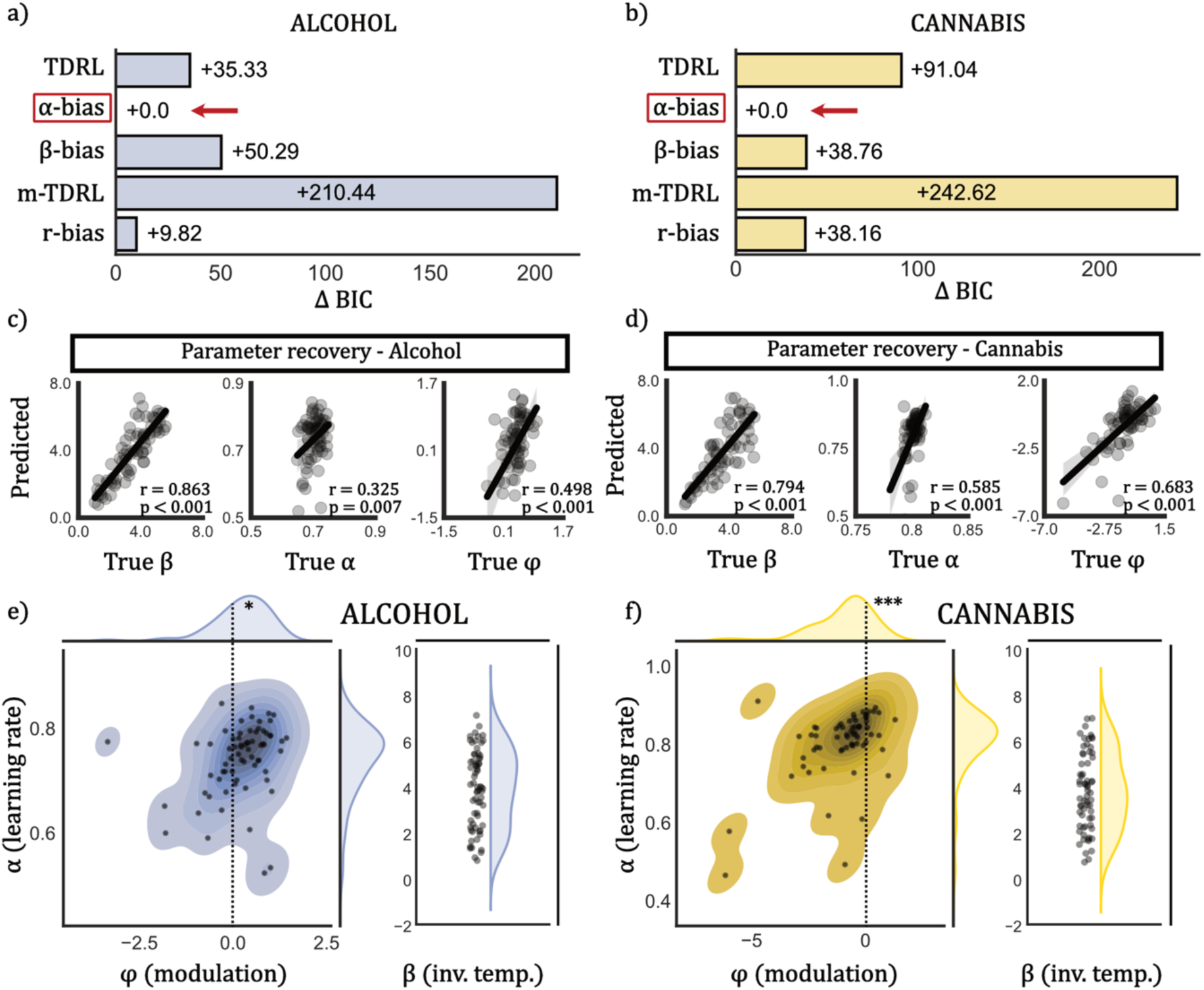
Decision-making model comparison and parameter distribution: addictive condition. *TDRL*: temporal difference reinforcement learning, *α-bias*: learning rate bias, *β-bias*: inverse temperature bias, *m-TDRL*: momentum-based TDRL, *r-bias*: reward bias. **(a, b)** For all models, ΔBIC was defined as the difference between each model’s BIC and the best performing BIC. The α*-*bias model performed best across groups. **(c, d)** For the best performing α-bias model, we performed parameter recovery by simulating data from parameter estimate, and refitting the simulated data. All parameters (α, β, φ) displayed excellent parameter recovery across groups (P<0.01 across parameters and groups). **(e, f)** Distributions of parameters were extracted from the α-bias model. The left panel displays the joint distributions of α (learning rate) and φ (modulation factor), as these interact directly in the model, while β (inverse temperature), is visualized separately. In the alcohol group, φ was found to be significantly positive (*t*=2.159, P=0.034), while in the cannabis group, it was found to be significantly negative (*t*=-5.590, P<0.001).

In the monetary condition, in contrast, the r-bias model performed best across groups (**Supp. Fig. 5a, b**) and parameter recovery for this model was, again, excellent (**Supp. Fig. 5c, d**). Examination of parameter values (**Supp. Fig. 5e, f**) revealed that both groups showed significantly positive φ (alcohol: M=0.118, SD=0.099, *P*<0.001; cannabis: M=0.190, SD=0.097, *P*<0.001), indicating that, across groups, higher craving levels increased the perceived magnitudes of monetary rewards.

These findings highlight an important role for craving in modulating learning across alcohol and cannabis groups. First, momentary craving biases *learning rate* in response to addictive cues, yet influences *reward perception* in response to non-addictive cues across both groups. Second, alcohol craving and cannabis craving have opposing effects on drug-related learning; alcohol craving accelerates alcohol-related prediction error encoding, while cannabis craving reduces learning based on cannabis-related prediction errors. These models provide overlapping yet distinct computational mechanisms mediating the relationship between craving and decision-making in alcohol and cannabis users.

### Trial-wise expectations and prediction errors combine to drive perceived craving across groups and decision contexts

Though our results corroborate a directional effect of craving on learning, the nature of the reverse interaction remains unclear, i.e., do how do prior expectations and outcomes influence perceived craving? Systematic model comparison revealed that, across both alcohol and cannabis groups, momentary craving was best explained by a combination of expected values and prediction errors in response to outcomes in both addictive and monetary conditions (**Fig. 3a, b**), rather than either individually. Predicted cravings generated by this model were also significantly correlated with true cravings (**Fig. 3c-f**; t>11.0, *P*<0.001). Overall, these results build on several recent findings substantiating the importance of both cue-induced and belief-induced influences on momentary craving^16,30^.

**Fig. 3.**
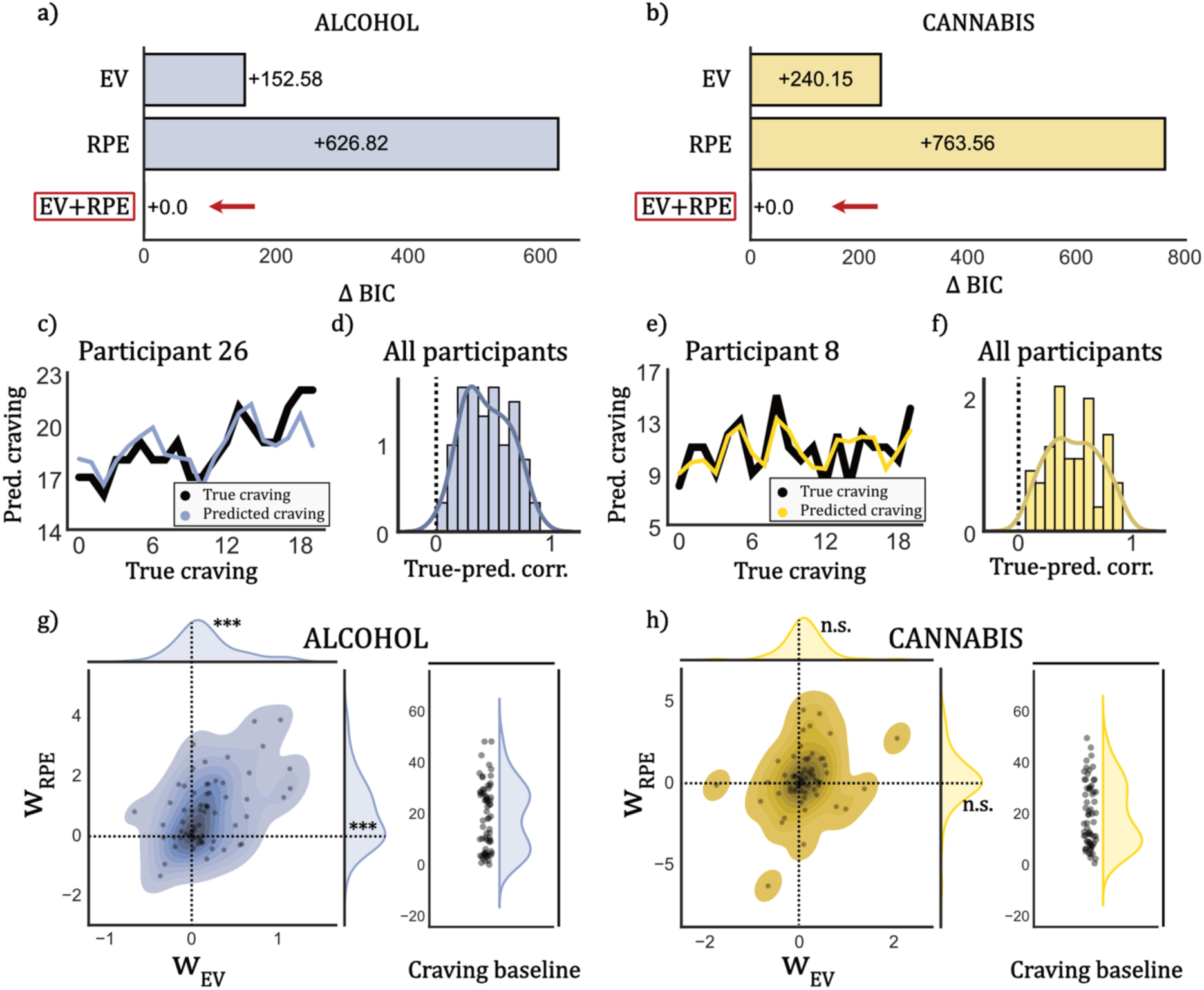
Craving model comparison and parameter distribution: addictive condition. *EV*: Expected-value-only craving model, *RPE*: reward-prediction-error-only craving model, *EV+RPE*: combined EV and RPE craving model. **(a, b)** As with decision models, ΔBIC was defined as the difference between each model’s BIC and the best performing BIC. The joint EV+RPE model performed best across groups. **(c-f)** In each group, for the best performing craving model, we calculated the correlations between model-predicted craving and true craving ratings. An example participant’s true vs. predicted craving are displayed in panels **c** and **e**. There was a high degree of correlation across participants (panels **d** and **f**; alcohol: mean *r*=0.438; cannabis: mean *r*=0.487), indicating strong model efficacy. **(g, h)** Distributions of parameters were extracted from the EV+RPE model. The left panel displays the joint distributions of *w_EV_* (EV weight) and *w_RPE_* (RPE weight), while *Craving baseline* is visualized separately. In the alcohol group, both *w_EV_* (*t*=3.983, P<0.001) and *w_RPE_* (*t*=4.940, P<0.001) were found to be significantly positive, while in the cannabis group, neither reached significance (*w_EV_t*=1.759, P=0.083; *w_RPE_t*=0.343, P=0.733).

Next, we extracted the parameters from the best performing model to interpret the processes underlying elicitation of craving during the task. In the addictive cue condition, EV weight was significantly positive in alcohol users (**Fig. 3g**; M=0.173, SD=0.358, *P*<0.001) but not cannabis users (**Fig. 3h**; M=0.108, SD=0.496, *P*=0.083. RPE weight was also significantly positive only in the alcohol users (M=0.664, SD=1.108, *P*<0.001), but not cannabis users (M=0.071, SD=1.680, *P*=0.733).

In the monetary condition, the combined EV+RPE model was again found to be best performing (**Supp. Fig. 6a, b**) and predicted cravings were highly correlated with true cravings (**Supp. Fig. 6c-f**). Analysis of parameter estimates (**Supp. Fig. 6e-f**) showed that EV weight was significantly positive in the cannabis users (M=0.162, SD=0.347, *P*<0.001), but not in alcohol users (M=0.010, SD=0.497, *P*=0.873), while RPE weight was non-significant for both groups (alcohol: M=0.219, SD=1.754, *P*=0.306; cannabis: M=0.317, SD=1.935, *P*=0.191).

In sum, the models constructed here provide a means for disentangling two important components of momentary perceived craving: effects of prior expectations (EVs) and effects of prediction errors (RPEs). Here, we again find highly divergent computational signatures for alcohol and cannabis users that are context-dependent. In the addictive cue condition, momentary craving is dynamically driven by increases in both EV and RPE for alcohol users, but not cannabis users. In the monetary condition, momentary craving is primarily driven by increases in EV for cannabis users but not alcohol users.

### Model-derived computational parameters have substance-dependent predictive utility

Thus far, our results provided a computational account for the bidirectional relationship between substance craving and decision-making. Next, we sought to examine if these computational estimates had utility in predicting clinical severity above and beyond simple demographics or model-agnostic metrics. Five classes of regressions were constructed: (1) Demographic regression (*Demo-only*), in which only basic demographics (age, sex, race, income, education level) were used to to predict severity, (2) Computational model-derived regression (*Comp-only*), in which only computational parameters from the addictive condition were used, (3) Model-agnostic regression (*Agnostic-only*), in which only task performance summary metrics (mean and s.d. of craving and choice optimality in the addictive condition) were used, (4) *Demo+comp*, where both demographic and computational predictors were included, and (5) *Demo+agnostic*, where both demographic and model-agnostic predictors were included.

For each group, models were compared and ranked by expected log pointwise predictive density Widely Applicable Information Criteria (elpd_waic) scores, and normalized true and predicted severity were plotted against each other (**Fig. 4a-d**). We found that the *Comp-only* model performed best in alcohol users (elpd_waic = -93.747; r=0.553, *P*<0.001), while *Demo-only* performed best in the cannabis users (elpd_waic = -94.188; r=0.373, *P*=0.002). We also sought to interpret the significantly predictive variables from the best-performing regression model in relationship to drug use severity scores (**Fig. 4e-f**). Alcohol use severity was positively associated with learning rate and baseline craving, and negatively associated with inverse temperature and EV weight. Cannabis use severity, however, was negatively associated only with age, education, and income.

**Fig. 4.**
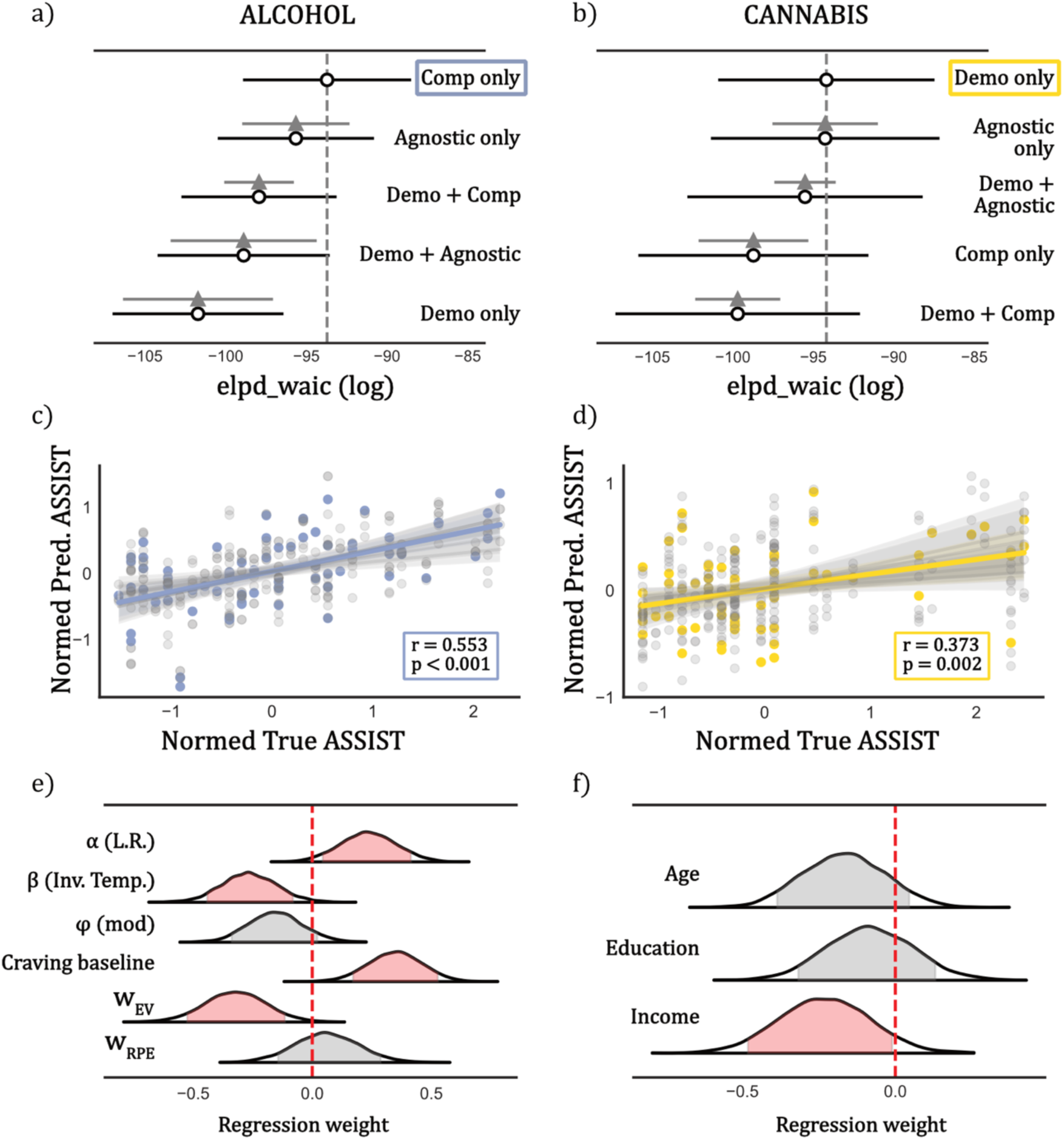
Clinical severity regression analysis. Regression analyses were conducted to determine the efficacy of computational parameters (*‘Comp only’*), model-agnostic metrics (*‘Agnostic only’*), demographics (*‘Demo only’*), and their combinations (*‘Demo+Comp’* and *‘Demo+Agnostic’*) in predicting clinical severity. Note that ASSIST scores were normalized prior to regression. **(a)** In the alcohol group, *Comp only* outperformed all alternatives (elpd_waic=-93.557). **(b)** In cannabis users, *Demo only* was the best performing model (elpd_waic=-94.032). **(c, d)** The best performing models in both groups generated predictions that were highly correlated with true ASSIST scores (alcohol: r=0.553, P<0.001; cannabis: r=0.373, P=0.002). **(e, f)** Parameters were extracted from the samples from the posterior distribution, and the 89% highest density intervals are plotted for each significant variable. Weight distributions for race and sex were eliminated due to high degree of variance and low contribution to prediction. In alcohol users, α (learning rate) and craving baseline were found to be significantly positively associated, while β (inverse temperature) and *w_EV_* were found to be significantly negatively associated. In the cannabis group, income was negatively associated, and there was a marginally negative association for age and education as well.

In sum, our comparative regression analysis unexpectedly found that computational parameters from our models were substance-dependent in their predictive utility. This may indicate that the direct utility of computational fingerprints of decision-making and craving in addictive disorders may be highly dependent on the addictive target and/or clinical group, or on a the particular set of computational latent parameters interrogated during the task.

## DISCUSSION

In the present study, we identified a computational mechanism subserving the dynamic relationship between momentary craving and decision-making generalizable across moderate-to-heavy alcohol drinkers and cannabis users. Our findings support the notion that craving and decision-making are two computationally intertwined processes across addictive domains. Additionally, we built on, and empirically tested, recent computational theories that prior expectations and prediction errors both play a role in perceived craving. Substantiating our hypotheses, we found that substance users converged on a generalizable algorithm in which momentary craving dynamically biased learning specifically in response to addictive cues, and both prediction errors and expected values influenced subsequent perceived craving. Notably, however, substance use groups diverged with respect to the patterns of parameters associated with decision-making and craving, providing distinct substance-specific computational fingerprints of these interacting mental processes in addiction.

There is compelling evidence supporting the view that computational mechanisms involved in addiction may be shared across substance use disorders, including alcohol and cannabis use. For instance, previous studies have highlighted several mechanisms for impaired goal-directed planning, belief-updating, and habit formation across substance use groups^41,42,46,47^. Network analytic approaches show craving plays a central role across substance use disorders^48–50^, and individuals who co-use alcohol and cannabis exhibit heightened cue-induced craving and altered decision-making for both substances^51–53^. Neurobiologically, several connectome-based predictive approaches have identified a transdiagnostic “craving network” involving regions of the salience, subcortical, and default mode networks^54–56^, suggesting a common neural signature for craving across addictive disorders, including alcohol and cannabis use disorders. Building on these prior results, our models provide first evidence that both alcohol and cannabis users converge on the same computational algorithm that explicitly links craving and decision-making.

Importantly, however, alcohol and cannabis users diverge in, and thus are uniquely identifiable by, their cognitive parameter patterns. In alcohol drinkers, we found that increased craving led to *faster learning* from alcohol-related prediction errors, suggesting that alcohol-associated outcomes that follow a state of high craving may lead to stronger valuation by the brain of a drinker. Several previous studies have identified similar learning dysfunctions in alcohol use disorder, including premature switching^57^, accelerated valuation of negative outcomes^58^, and high impulsivity^59^ that may be reflective of faster craving-induced learning. In contrast, in cannabis users, high craving led to *slower learning* from prediction errors about cannabis related outcomes. This result conforms with previous findings of diminished learning in cannabis users^60^, particularly as relating to memory^61,62^ and social context^63^ of cannabis cues. Additionally, while direct comparisons are lacking, several studies have also described clear dissimilarities in computational factors^64,65^ between alcohol and cannabis users (e.g., impaired delay discounting but unaffected reversal learning in cannabis users, but significant reversal learning impairment in alcohol users). Differences in computational signatures may also reflect the the disparities in clinical phenomenology between alcohol and cannabis consumption; for example, alcohol is often consumed in shorter, concentrated time periods, and its intoxicating effects take longer than those of cannabis, which is often consumed more gradually throughout longer periods of time and takes only minutes for its effects to be felt. Overall, our findings propose drastically diverging alterations in the craving-learning associations present in these two substance use disorders that, in turn, might guide development of substance-specific clinical interventions. Further exploration and characterization will be necessary to validate these findings prospectively, explore brain substrates, and compare/contrast them to the computational signatures of other substance use disorders (e.g., opioid, cocaine).

We also found that, across groups, both *expected value* and *prediction error* signals – regardless of the decision context – dynamically drove changes in cravings. Importantly, this finding realigns the classic cue-induced craving phenomenon^9,66^ with well-known neural^67,68^ (dopaminergic) and computational (prediction error)^16,32,44^ effects. In this view, cue-reactivity can be deconstructed as being composed of both momentary prediction errors and expected values that each independently drive increased craving; i.e., cues elicit craving through two distinct mechanisms: (1) prior beliefs about their value (EV-driven effect), and (2) epistemological surprise (RPE-driven effect). This result explains recent findings that both values and prediction errors are important components of perceived craving in addiction at behavioral^69–72^, computational^41,73,74^ and neural^16,75,76^ levels.

Finally, we found that, among alcohol users, computational parameters can reflect latent information that is predictive of clinical severity above and beyond model-agnostic metrics or demographics, suggesting that these latent computational parameters may contain unique information related to individual differences in alcohol use severity. To our knowledge, this provides the first empirical evidence that computationally-derived parameters may capture features core to addiction that are not provided by model-agnostic measures. These findings provide compelling evidence for the value of computational modeling in uncovering latent information that may be highly valuable translationally. Nevertheless, further work will be required to refine the sets of latent computational parameters that may be most salient to the predictions of clinical severity in other addictive disorders.

Our finding of an algorithmic link between craving and decision-making has direct clinical implications: specifically, by reducing or managing cravings, individuals may be able to break the vicious cycle of addiction-associated decision-making that leads to poor clinical outcomes. Though interventions regarding craving have been proposed, tested, and examined widely^77–79^, our models provide the first mechanistic explaination for their success. Further exploration may reveal a novel line of therapeutic interventions focusing on the interplay between craving and addiction-associated decision-making that target both components simultaneously. Though recent psychological research has hinted at similar results^71,80,81^, this finding certainly merits further clinical investigation, as the majority of current interventions typically involve focusing on negative aspects of addiction directly^72,82,83^, or mindfulness strategies^84–86^.

Our results here are limited by an absence of direct evidence supporting the neurobiological substrates that facilitate this mental computations proposed in this model. Though numerous efforts have identified the effect of decision-making on subjective states, such as happiness^28^ or craving^16,75^, there are, as of yet, no neuroimaging studies investigating a bidirectional interaction between the two. A next logical step would be to collect neural data (e.g., using neuroimaging) in conjunction with our experimental design, and utilize machine learning or statistical approaches to find the neural patterns encoding the computational mechanisms outlined here. Additionally, these findings would need to be replicated in a sample of individuals with confirmed substance use disorders to ensure the applicability of the model to true clinical populations, and allow for a more robust understanding of how the proposed mechanisms operate under pathological conditions.

## CONCLUSION

Overall, this study highlights a reciprocal, dynamic link between decision-making and craving in addictive disorders. Our results demonstrate that momentary craving leads to biased learning in drug-salient contingencies, and support an updated and refined craving framework, with both prior beliefs and prediction error effects being key drivers of momentary perceived craving.

## METHODS

### Task design - instructions

We utilized a modified two-armed bandit task that incorporated reversals to encourage continuous learning during the experiment. At the start of the task, participants were presented with the instructions about how to play the game. Participants were presented with two slot machines and were encouraged to do their best to maximize their rewards from the machines, with the incentive of being rewarded with a greater bonus payment at the end of the game based on their final score.

In the money condition, participants received a monetary reward, presented as an image of a coin. In the addictive reward condition, they received a reward corresponding to an addictive cue, selected based on the addictive group to which they belonged (e.g., in the alcohol-using group, participants were rewarded with an image of alcohol). Additionally, during the start of the experiment, participants were able to select one of three possible addictive cues that were most tempting to Participants in the alcohol-using group were able to select from either a beer, wine, or liquor reward, while cannabis users were able to select from either a blunt, bong, or bowl.

Participants were then also informed that they would intermittently be asked to assess and report their craving and mood. They were given specific definitions for craving (“*An intense, conscious desire or wanting for something*”) and mood (“*A non-specific, persistent general feeling about your current mental state, distinct from emotions, which are shorter-lived and specific to a particular thing*”), in order to provide them with a deeper understanding of the concepts being measured, and to provide a concrete basis upon which their self-reports could be compared. Baseline craving and mood ratings were then assessed prior to presentation of the slot machine task.

Finally, participants were informed that one of the slot machines would be more rewarding than the other at all times, but the more rewarding slot machine might change over the course of the experiment. The true experimental structure (delineated below), including the probabilities of reward, the reversal timings, and number of trials, was not revealed to the participants beyond the information listed above.

### Task design - experimental structure

There were 60 trials per condition, and 5 practice trials at the start of the experiment for a total of 125 trials. The 5 practice trials were discarded during the computational modeling. Presentation of conditions was randomized over the cohort, such that half of the participants received the money condition first and the other half received the addictive cue condition first. The more rewarding slot machine presented a reward 80% of the time, while the less rewarding machine presented a reward 20% of the time. The more rewarding machine switched four times over the course of the experiment (“reversals”). First selection of the best machine (either right or left), was pseudorandomized and reversal timings were also pseudorandomized as either 12-12-11 or 13-12-10. Craving ratings were assessed approximately every three trials for a total of 20 craving ratings per condition. Mood ratings were assessed approximately ever 5 trials for a total of 12 mood ratings. Ratings were collected on a linear scale from 0-50, where participants used the arrow keys to move a cursor from left to right to select their answers, where the left end of the bar was labeled as ‘Low’ and the right end was labeled as ‘High’.

### Data collection

We first performed a high-throughput screening for 1,000 participants on Prolific, an online data collection platform for recruiting participants across the world. We restricted our recruitment to participants with the following characteristics: USA residents, fluent English speakers, high approval rating on Prolific, no previous completion of any of our group’s experiments. Furthermore, we recruited a balanced sample of male and female participants. Participants completed two screening surveys implemented on Redcap. The first was the World Health Organization’s Alcohol, Smoking and Substance Involvement Screening Test (ASSIST)^87^, which was developed as a screening tool to help primary health professionals detect and manage substance use and related problems. ASSIST allows for screening of use of substances such as alcohol, cannabis, tobacco, stimulants, inhalants, and others. Second, we used a lab-standard demographics survey to assess basic demographic features such as age, sex, level of education, income, and race.

From the screening surveys, we recruited cohorts for moderate to high alcohol use and cannabis use. Potential participants were identified with the following criteria: (1) The alcohol use cohort reported moderate to heavy use of alcohol weekly as identified by ASSIST, with no comorbid usage of any other substances. (2) The cannabis use cohort reported moderate to heavy use of cannabis weekly, with low to no use of alcohol, and no other substance usage.

From the full eligible cohort for each group, we collected a total of 40 participants’ data for the slot machine game. Participants with substance use were additionally asked to complete further group-specific questionnaires. Alcohol users completed an Alcohol Dependence Scale^88,89^, and an Alcohol Use Disorder Identification Test^90^. Cannabis users completed a Cannabis Severity of Dependence Scale^91^, and a Cannabis Abuse Screening Test^92^.

We followed the exact protocol listed above for a second sample, starting with 1,000 screened participants, filtering of participants into potential cohorts for data collection, collection of 40 samples from each group of interest, and collection of auxiliary clinical questionnaires for the substance use groups. The data from the two samples was combined into our final analysis sample reported in the manuscript.

### Quality control

Before analysis, participant data from each group underwent a series of quality control checks. First, we included one attention check during the experiment, with instructions about halfway through the task to select the highest option if the participant was paying attention. Participants that did not pass this attention check were excluded. Second, since the optimal machines and reversal timings are known, it is possible to construct the vector of optimal choices during the experiment. A participant’s raw choice behaviors were compared to the optimal choice vectors for each condition, and only participants with greater than 50% optimality were selected, ensuring that they learned to play the game well, and were not simply randomly responding in the task, with a relatively low bar of exclusion reducing the chance that randomness of reward structure did not unnecessarily exclude well-performing participants. Finally, we z-scored the reported craving ratings during the task and excluded the participant if the standard deviation of the ratings did not exceed 1, suggesting that there was very low variability in the rating scores. Moreover, we found that most participants with extremely low craving variability seemed to exclusively report a craving of 25/50, which was the default value in the rating scale, suggesting that they were not properly responding to the prompts.

### Model-agnostic analysis

1. **Calculation of mean/SD of craving and mood:** We calculated the means and variances of reported cravings for each individual across groups and conditions. These were used to assess whether simple summary statistics of overall trends of craving were associated with clinical measures. The same process was repeated for reported moods. Individual distributions of cravings and moods were aggregated and reported across groups and conditions.
2. **Survey scoring:** We scored the Redcap surveys for each group according to a questionnaire-specific scoring. For the alcohol group, we utilized 3 surveys: ASSIST, AUDIT, and ADS. ASSIST scores ranged from 0-40 (<3 – low severity, 4-26 – medium severity, >26 high severity). AUDIT scores ranged from 0-34 (<7 – low severity, 8-14 – medium severity, >14 high severity). ADS scores ranged from 0-54 (0 – No risk, 1-13 – low risk, 14-21 – moderate risk, 22-30 – substantial risk, >30 – severe risk). For the cannabis group, utilized 3 surveys: ASSIST, CAST, and SDS. ASSIST-Cannabis scores ranged from 0-36. CAST scores ranged from -6-20 (<3 – low risk, 3-6 – moderate risk, >6 high risk). SDS scores ranged from -5-13 (<=3 – low risk, >4 – high risk). The distributions of all these surveys were visualized across groups. Additionally, we calculated the difference between clinical severity scores before the task (during screening surveys) and after the task to assess the stability of these clinical measures.
3. **Choice optimality:** We qualitatively assessed the performance of participants to ensure that they were able to learn the structure of the task well. To do this, we compared the choice of machine for all participants in a group to the optimal choice that could be made at that time, where the optimal choice was defined as the machine with higher probability of reward. Participants with lower than 50% optimality were excluded from further analysis because they were unable to learn the task structure. Finally, we plotted the distributions of optimality of remaining participants to demonstrate that participant optimality was well above chance.
4. **Qualitative performance checks:** We averaged the choice of machine across participants by group, controlled for randomization of the order of the best machine presented first (either left or right) to qualitatively assess group-level tracking of reversals across the task. We then visualized the overall distributions of craving and mood during the task within individuals and across groups to ensure that there was a realistic distribution of cravings and moods reported during the task.
5. **Low-level sanity check correlations:** We correlated group-specific clinical scores (ASSIST for alcohol and cannabis, EDEQ for binge-eating, and usage survey for social media) with percent optimality, total score, and baseline and mean craving/mood ratings. We also performed a Spearman correlation between reported mood and craving ratings within participant to check the covariance of the two.

### Modeling analysis

Decision-making and craving modeling was done sequentially. There were five classes of decision-making balanced on model complexity (i.e., number of parameters utilized by the model).

1. Asymmetric temporal difference learning – Values were computed using a standard temporal difference reinforcement learning (TDRL) rule with asymmetric learning from positive and negative prediction errors^39,93^. Decisions were made with a softmax rule.

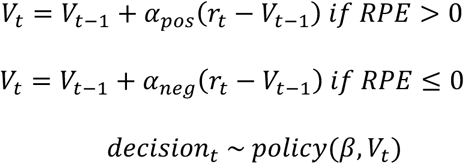
2. Reward modulation models – Values were computed with TDRL but reward magnitude was modulated by a bias parametrized by momentary craving and a modulation factor *φ*^41,42^. Decisions were made with a softmax rule.

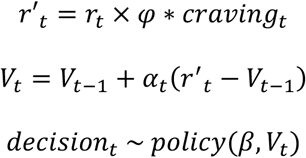
3. Learning bias models (*α bias*) – Values were computed with TDRL but learning rate *α* was modulated by a bias parametrized by momentary craving and a modulation factor *φ*. Decisions were made with a softmax rule.

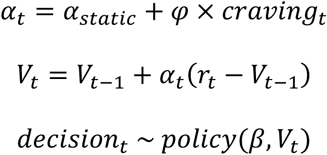
4. Temperature bias (*β bias*) – Values were computed with standard TDRL but inverse temperature *β* was modulated by a bias parametrized by momentary craving and a modulation factor *φ*. Decisions were made with a softmax rule.

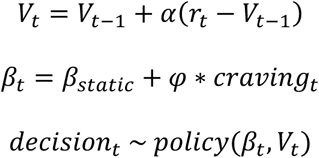
5. Momentum-based model (m-TDRL) – Values were computed with TDRL but reward was modulated by a momentum term parameterized by nonlinear effects of past prediction errors^24,43^. Decisions were made with a softmax rule.

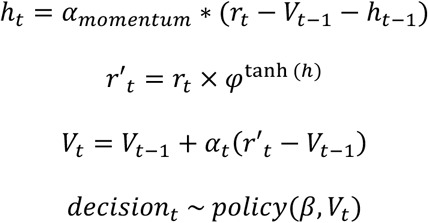

Following decision modeling, momentary craving was modeled as a non-linear combination of prior beliefs (i.e., expected values) and momentary surprise (i.e., prediction errors). There were three variants of craving models:

1. Prediction-error-elicited craving (*RPE*) - In this class, momentary craving was modeled as the geometrically decaying effect of momentary prediction errors, along with a static baseline craving.

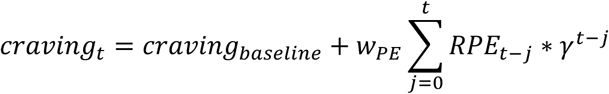
2. Expectation-elicited craving (*EV*) - In this class, momentary craving was modeled as the geometrically decaying effect of momentary expected values, along with a static baseline craving.

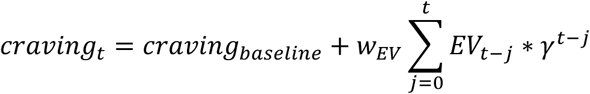
3. Jointly elicited craving (*EV+RPE*) - In this class, momentary craving was modeled as the geometrically decaying effect of both momentary expected values and momentary prediction errors, along with a static baseline craving.

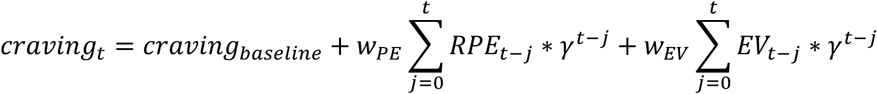

### Model implementation

Decision and craving models were implemented using pyEM^94–96^, a Python library for parameter estimation with iterative expectation maximization. Choices and craving ratings were modeled independently in two stages. During the expectation step, the maximum likelihood probability (P_MLE_) of task choices for each participant *i* was defined as the conditional probabilities of each choice at trial *t* given the expected values (EV_t_) for both slot machines at that trial and the participant’s parameter vector θ_i_ (i.e. ∑ log (*p*(choice_t_|EV_t_, θ_i_))). The prior probability (P_prior_) was defined as the log-likelihood of the participant’s θ_i_, given the current group–level Gaussian prior distributions of the parameters (**θ**) across participants, with a mean vector **μ** and standard deviation **σ*^2^***. The maximum a posteriori (MAP) estimate was calculated by maximizing the sum of log P_MLE_ + log P_prior_ across participants. Subsequently, during the maximization step, the group-level Gaussian prior distributions (parametrized by **μ** and **σ*^2^***) were recomputed given the **θ** vectors computed in the prior expectation step. These steps were repeated until convergence, where MAP changed by <0.001 in consecutive iterations, or a maximum of 800 steps. The geometric decay parameter γ was estimated freely in the first iteration, and then fixed to the mean of the estimates in the second iteration of model fitting in order to aid with model convergence. The group level priors were initialized at **μ**=0.1 and **σ*^2^***=100 in order to allow for uninformative data-driven priors. Decision model parameters were transformed from the Gaussian parameter space to the decision-model space by applying the appropriate link functions. A sigmoid function was applied to the learning rate (α). Inverse temperature was defined as *β_transform_* = 10⁄1 + *e^−β^*. The exact same procedure was applied to the craving ratings, except that trial-wise log-likelihood value was defined as conditional probability of the observed craving rating from a Gaussian centered at the predicted model value at that trial.

### Model fit metrics

Posterior samples of parameter sets were used to simulate choices for each participant following parameter fitting. The model ran a simulation for each parameter set for a total of 4,000 datasets of simulated choices. The following checks were performed on the simulated data.

1. Means of these simulations, were visualized against true actions to qualitatively assess congruence.
2. Each choice simulation was given an accuracy score representing the number of simulated actions that matched true actions, where 50% was chance accuracy.

### Model comparison

Models were compared using the integrated Bayesian information criterion (iBIC) score^96,97^. Briefly, the iBIC score was calculated with the following steps. *k*=2000 samples were drawn from the final group-level Gaussian estimates for parameters **θ**(**μ**, **σ**^2^) for each participant *i*. The log-likelihood of each sample (LL_i,k_) was computed as the sum of the conditional probabilities of the participant’s choices (choices_i_) given the sample parameter vector θ_k_. This value was calculated for each sample, and across participants. LL_i,k_ were then summed across all participants and samples with the following equation: *iLog* = ∑*_i_* log (∑*e^LLi,k^*/2000), and iBIC was defined as *iBIC* = −2 × *iLog* + *n_param_* × log (*n_trials_*). This procedure was applied to each model independently. The model with the lowest score was defined as the reference model, and ΔBIC scores were calculated for each model as the difference from reference model iBIC.

### Parameter estimate distributions

Across groups, and within each condition (money or addictive cue) and model type (decision or craving), we identified the best performing model by the ΔBIC score. For this model, we plotted the distributions of parameter estimates across participants for all parameters utilized in the model. We visualized decision-making parameters and craving parameters in separate sub-figures for easier summarization and interpretation. For each parameter, we tested the directionality of the estimated effect by calculating statistical significance from zero. All significance testing was performed with parametric t-test (either independent or relative, depending on suitability of the samples) and confirmed with non-parametric Mann-Whitney U-tests.

### Clinical score prediction

We used a multiple linear regression model to test the hypothesis that joint parameter estimates from best performing models can successfully predict clinical severity scores. For clinical severity scores, we decided to use ASSIST, a group-specific survey that had high variability across participants. The best performing decision-making and craving models identified by the model comparison procedure, restricted to the more salient addictive cue condition, was tested in five classes of regression analysis. In the demographic regression (Demo only), the participant-specific demographic information (i.e., age, sex, education level, self-reported race, and income) were used as regressors. In computational regression (Comp only), decision- and craving-model parameter estimates were used as regressors. In model-agnostic regression (Agnostic only), simple means and variances of craving and choice optimality were used to predict clinical severity. In the last two models, demographic regressors were combined with computational regressors (Demo + Comp) and agnostic regressors (Demo + Agnostic) respectively. Models were implemented using ‘bambi’, a Bayesian linear modeling Python library^98^. Final reported models represent the best-performing (by AIC and ELPD-WAIC) models. Variables not contributing significantly to prediction, demonstrated by large variation is weight estimates, were omitted (e.g., race and sex in the cannabis group).

**Supp. Fig. 1.**
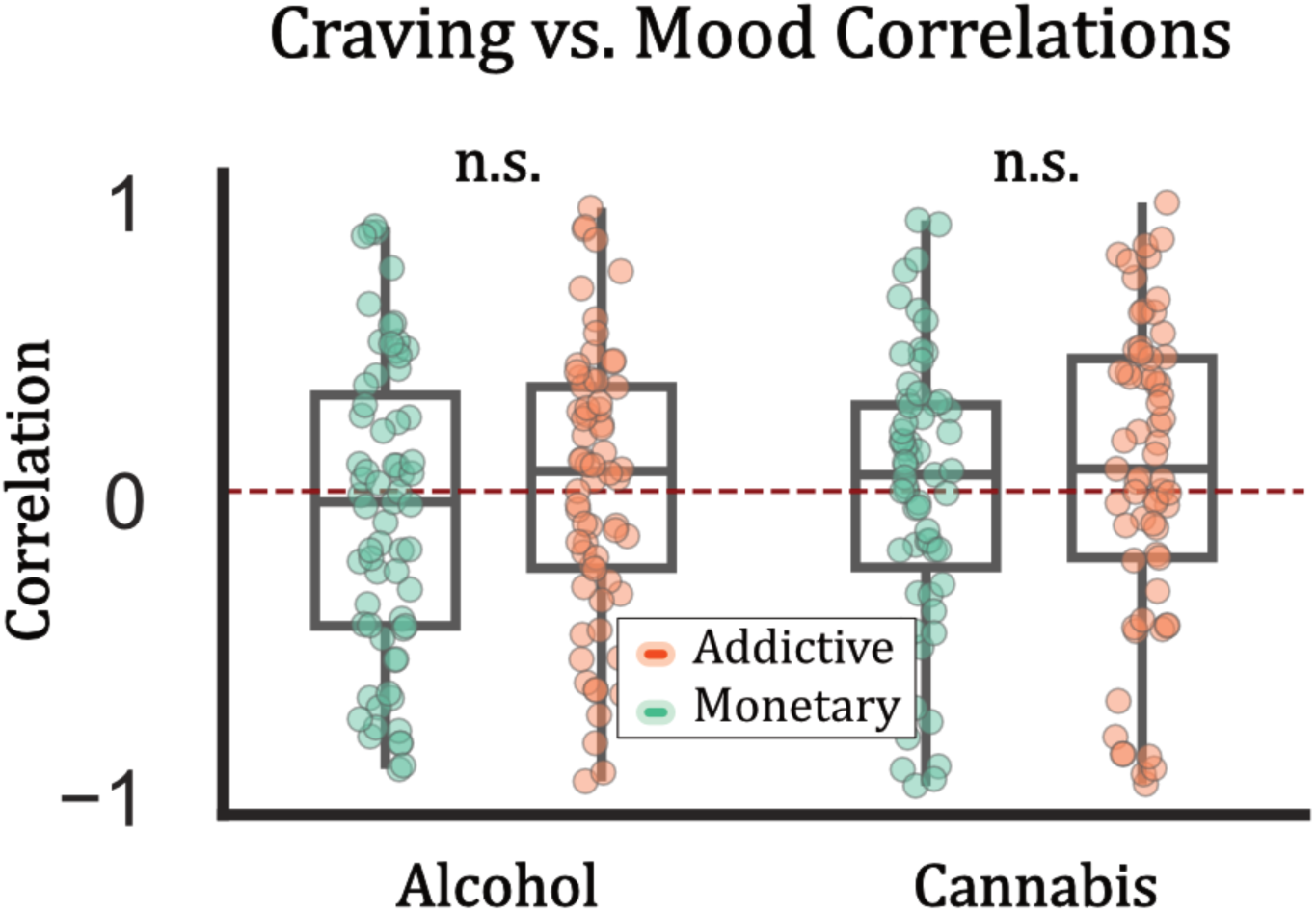
Correlations between craving ratings and mood ratings. Within each group, there was no significant relationship between craving and mood ratings within or across conditions. For the alcohol group, mean correlation was r=-0.063 (P=0.305) for the money condition, r=0.021 (P=0.703) for the addictive condition, and the difference between conditions was not significant (*t*=1.098, P=0.276). For the cannabis group, mean correlation was r=-0.002 (P=0.968) for the money condition, r=0.065 (P=0.308) for the addictive condition, and the difference between conditions was not significant (*t*=0.972, P=0.334).

**Supp. Fig. 2.**
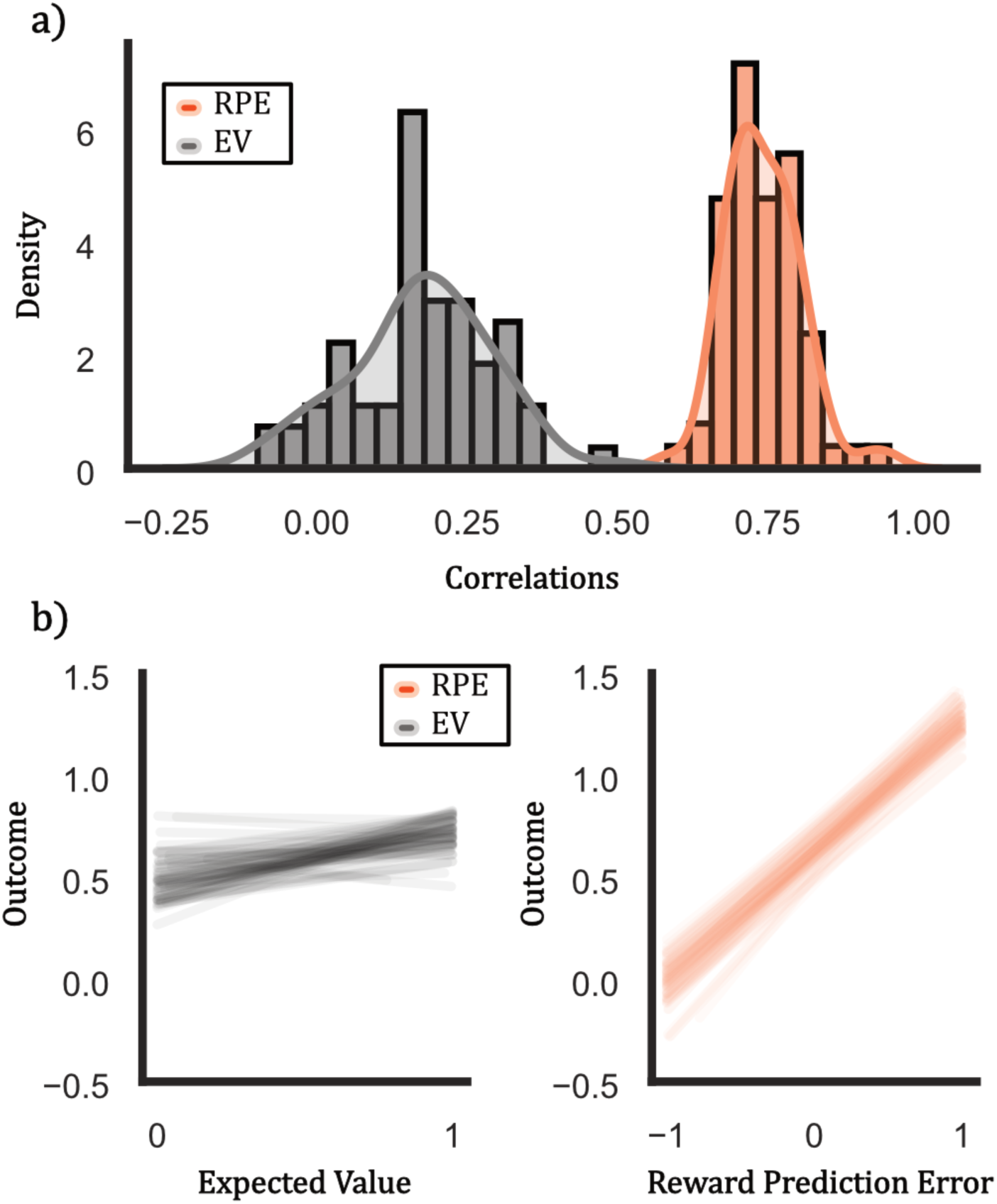
Correlation between RPE/EV and outcomes. To validate the use of reward prediction error (RPE) based craving models as a proxy for cue-induced craving, we calculated the correlations between trialwise RPE and outcomes. Intuitively, note that every positive outcome (‘win’, presentation of cue) will result in a positive RPE, while every negative outcome (‘loss’, absence of cue) will result in a negative RPE. This relationship would not necessarily be present for the association between expected value (EV) and outcome. (a) On average, participant-wise correlation between RPE and outcome was M=0.742, SD=0.062, while correlation between EV and outcome was M=0.174, SD=0.116. (b) For all participants, the line of best fit is shown between EV vs. outcome (left, gray), and RPE vs. outcome (right, orange).

**Supp. Fig. 3.**
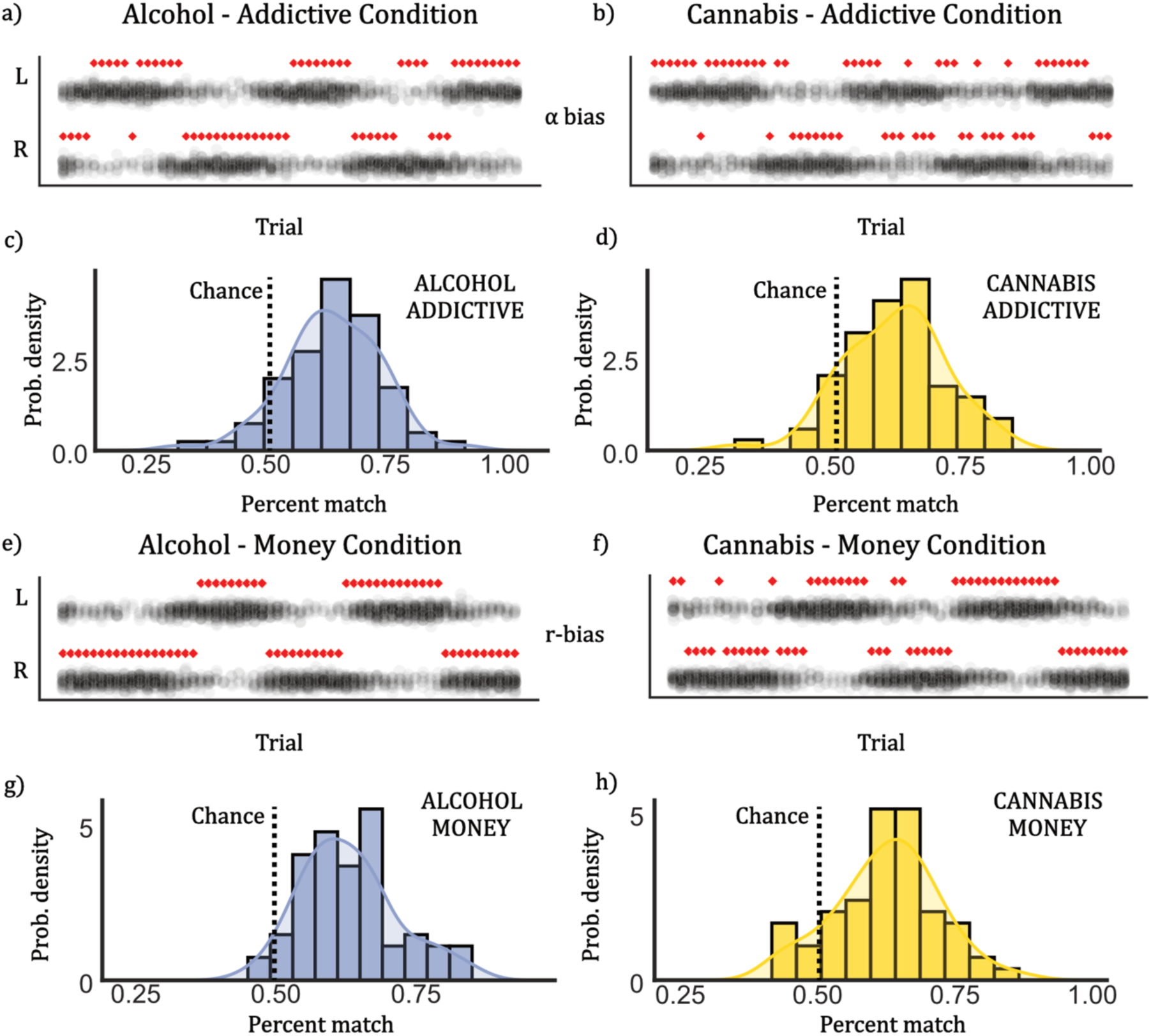
Decision-making model simulation. Simulated choices were generated from candidate models after model fitting. 4000 simulated datasets were generated and compared to true choice behavior on a session-by-session basis. (a, b) 50 sets of decisions for a sample participant in the addictive condition are plotted as black circles, and qualitatively have a high similarity to true choices (red diamonds). (c, d) We quantified the percent of matches in true and simulated choices, and statistically compared to chance (50%). Simulations matched true behaviors significantly better than chance (alcohol: *t*=12.405, P<0.001; cannabis: *t*=13.932, P<0.001). (e-h) We repeated the same procedure in the money condition with similar qualitative and quantitative findings (alcohol: *t*=10.198, P<0.001; cannabis: *t*=11.589, P<0.001)

**Supp. Fig. 4.**
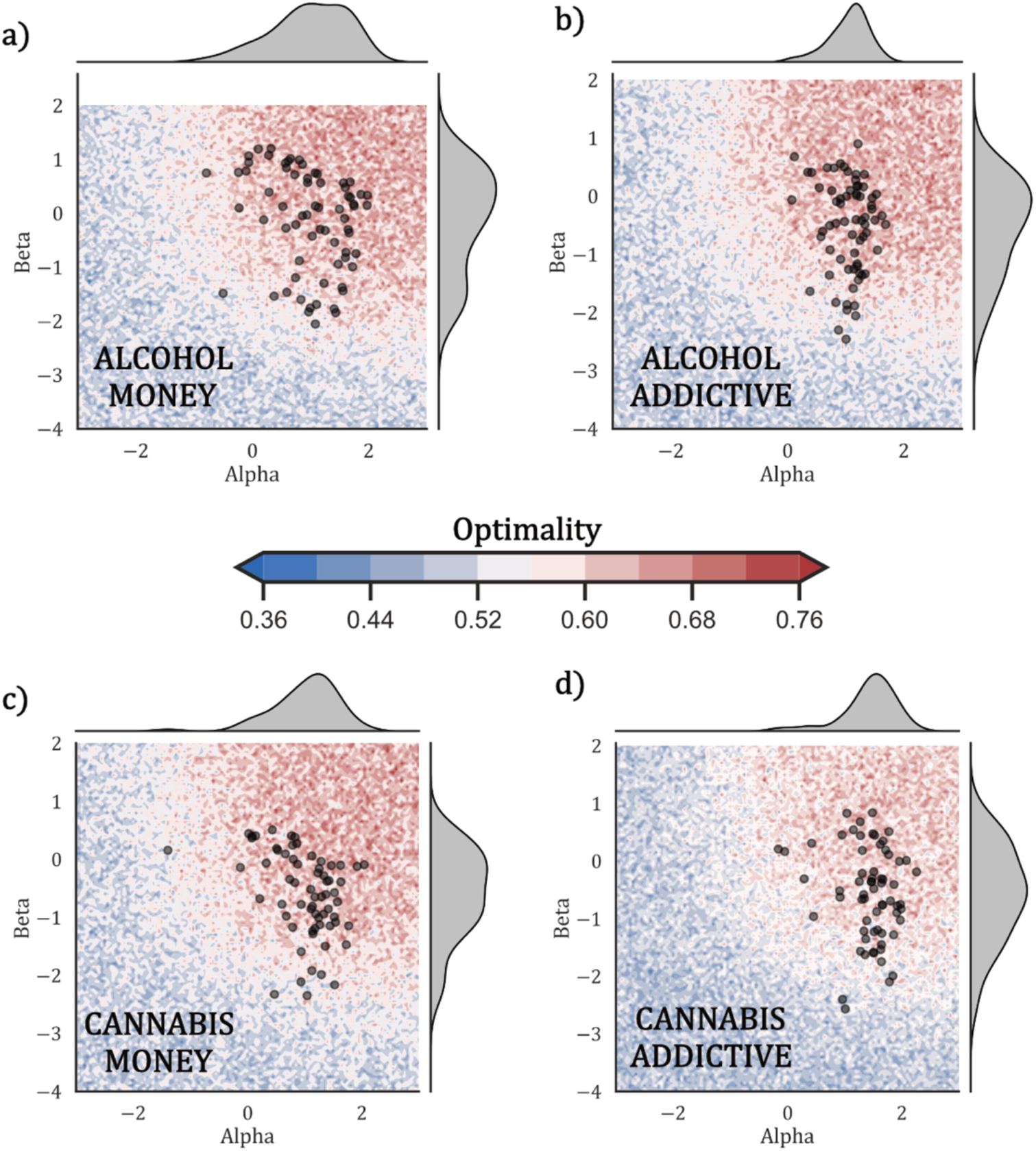
True vs. optimal parameter distributions. To confirm our simulation findings that participants behaved close to optimally, we performed a baseline simulation, randomly sampling (untransformed) α (learning rate) and β (inverse temperature) to discover the range of parameters leading to optimal outcomes (measured as percent correspondence with true task structure). Note that φ (modulation factor) is not included for this simulation. For each group-condition pairing, the joint parameter space is shown in **(a-d)** with red signifying highest optimality. Participant parameter estimates derived from the best performing models are overlaid as black dots. As expected, the large majority of participant estimate lie within the optimal range of parameters.

**Supp. Fig. 5.**
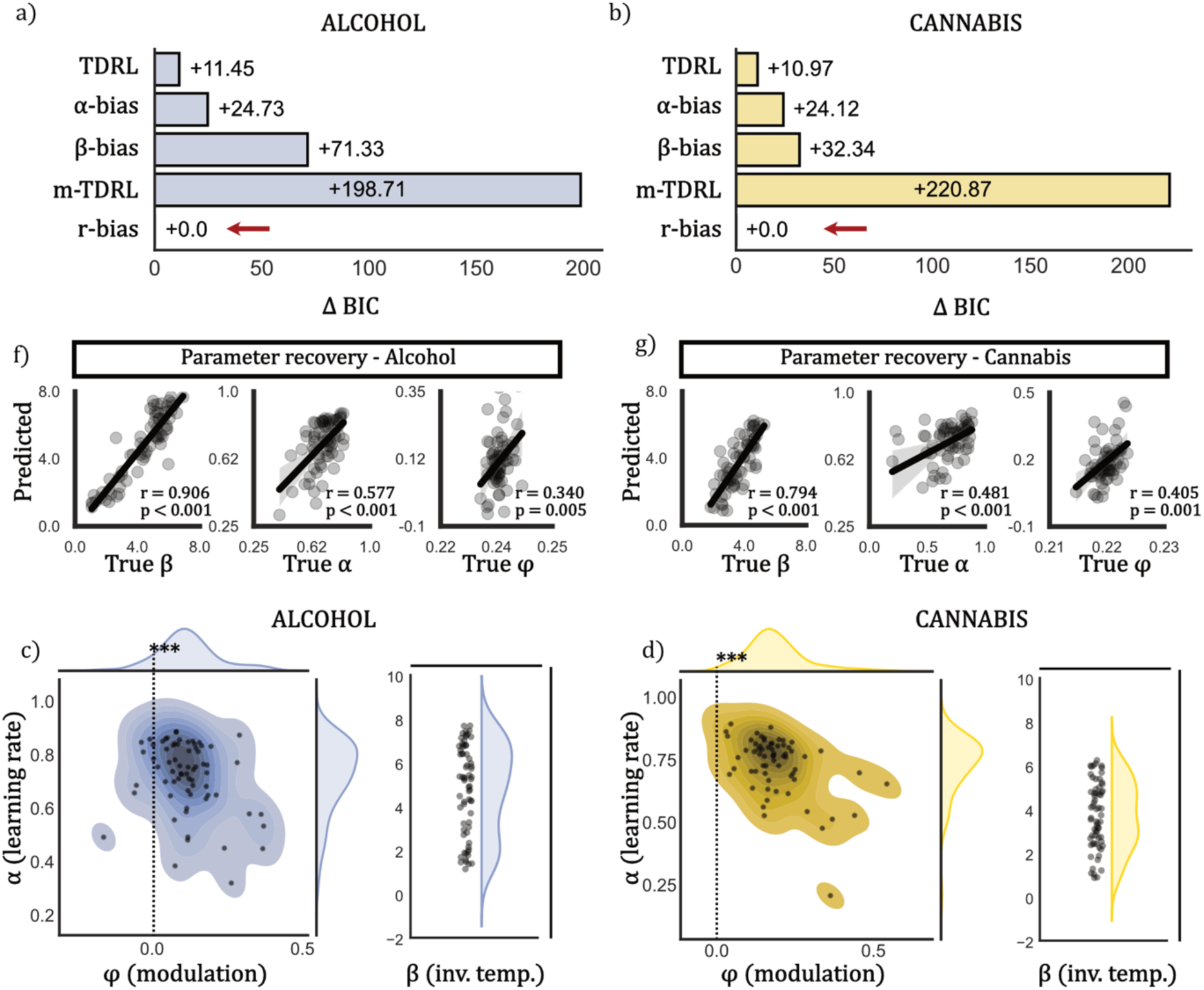
Decision-making model comparison and parameter distribution: money condition. **(a, b)** As with the addictive condition, ΔBIC was defined as the difference between each model’s BIC and the best performing BIC. The r-bias (reward bias) model performed best across groups. **(c, d)** All parameters (α, β, φ) displayed excellent parameter recovery across groups (P<0.01 across parameters and groups). **(e, f)** Distributions of parameters were extracted from the r-bias model. The left panel displays the joint distributions of α (learning rate) and φ (modulation factor), as these interact directly in the model, while β (inverse temperature), is visualized separately. In both the alcohol group and the cannabis group, φ was found to be significantly positive (alcohol: *t*=9.812, P<0.001; cannabis: *t*=15.831, P<0.001)

**Supp. Fig. 6.**
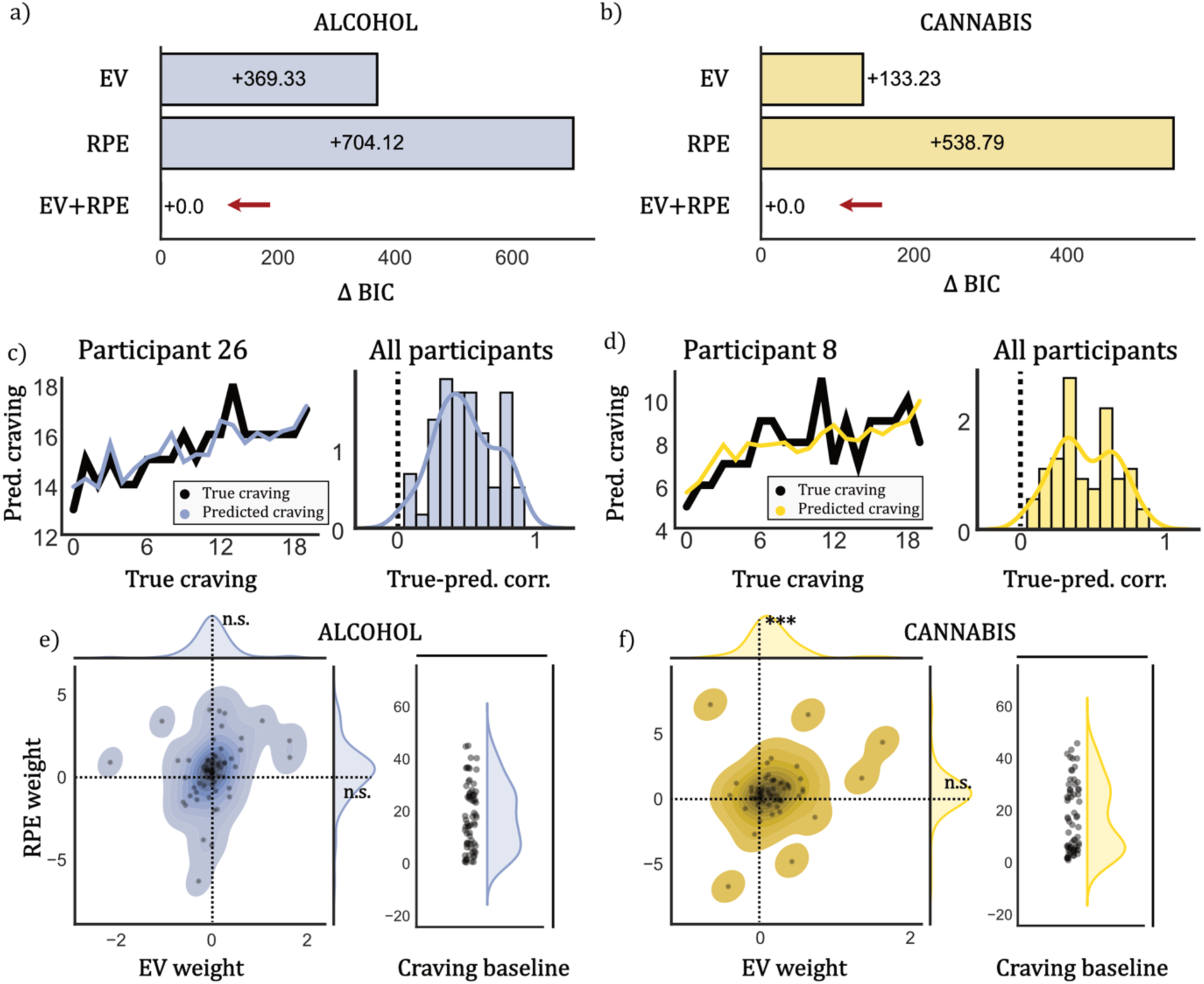
Craving model comparison and parameter distribution: money condition. **(a, b)** Again, ΔBIC was defined as the difference between each model’s BIC and the best performing BIC. The joint EV+RPE model performed best across groups. **(c-f)** In each group, for the best performing craving model, we calculated the correlations between model-predicted craving and true craving ratings. An example participant’s true vs. predicted craving are displayed in panels **c** and **e**. There was a high degree of correlation across participants (panels **d** and **f**; alcohol: mean *r*=0.489; cannabis: mean *r*=0.446), indicating strong model efficacy. **(g, h)** Distributions of parameters were extracted from the EV+RPE model. The left panel displays the joint distributions of *w_EV_* (EV weight) and *w_RPE_* (RPE weight), while *Craving baseline* is visualized separately. In the alcohol group, neither *w_EV_* (*t*=0.160, P=0.873) nor *w_RPE_* (*t*=1.031, P=0.306) reached significance, while in the cannabis group, *w_EV_* was positively associated (*t*=3.765, P<0.001), but *w_RPE_* was not (*t*=1.321, P=0.191).

**Supp. Fig. 7.**
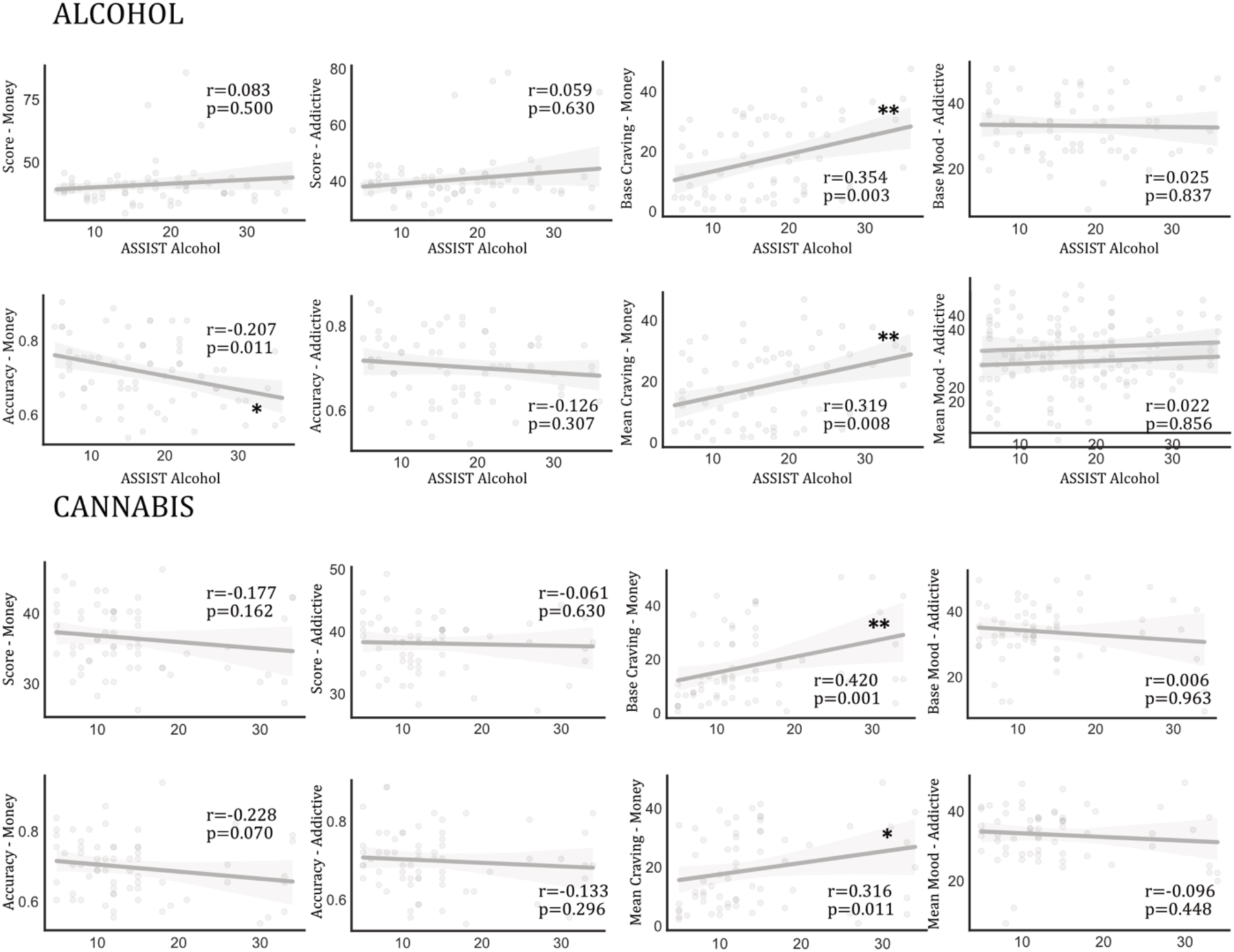
Correlations between model agnostic task performance, craving ratings and mood ratings. We correlated mean and initial (base) craving and mood, as well as choice optimality and task score with clinical severity scores directly. We also discovered an expected positive relationship between alcohol/cannabis use severity and both mean and initial craving ratings (r>0.319, P<0.008), but no relationship existed between mood and severity scores (r<0.096, P>0.448). There was also a negative correlation between clinical severity score and accuracy in the money condition for alcohol users (r=-0.207, P=0.011).

